# Critical amino acid residues in the N-terminal domain of NADPH-dependent assimilatory sulfite reductase flavoprotein mediate octameric assembly

**DOI:** 10.64898/2026.04.08.717228

**Authors:** Nidhi Walia, Thais Pedrete, Fozieh Ahmadizadeh, Ekhtiar Rahman, Yashika Garg, Brian Washburn, Cheryl Pye, Fanny C. Liu, Peter Randolph, Kevin L. Weiss, Gergely Nagy, Christian Bleiholder, M. Elizabeth Stroupe

## Abstract

How large, flexible enzymes assemble into defined oligomeric architectures remains a central question in biology. NADPH-dependent assimilatory sulfite reductase (SiR) forms a heterododecamer built on an octameric flavoprotein (SiRFP) core, yet the molecular basis for this assembly has been unresolved because of its disordered N-terminus. Here, we use ion mobility mass spectrometry, small-angle neutron scattering, and mutagenesis to define the mechanism of SiRFP oligomerization. We show that SiRFP forms a discrete, stable octamer in solution. We also report that its N-terminal 52-residue segment is necessary and sufficient to mediate assembly, also mediating oligomerization when fused to a heterologous protein. Structure-guided mutagenesis identifies four residues (Gln22, Tyr39, Phe40, and Gln47) whose substitution disrupts the octamer, producing concentration-dependent lower-order species while retaining catalytic activity. These findings define the determinants of SiRFP assembly with broader implications for engineering homomeric protein complexes.

**Importance:** This work seeks to understand the basis for oligomerization of a large oxidoreductase that is important for metabolizing sulfur, an essential chemical for all of biology. A 52-residue long leader peptide is necessary and sufficient for assembly into a particularly stable octamer that is resistant to chemical denaturation under diverse conditions.

## Introduction

NADPH-dependent sulfite reductase (SiR) plays a crucial role in proteobacteria by converting sulfite (SO_3_^2-^) to sulfide (S^2-^) (Siegel *et al*., 1973), the most bioavailable form of sulfur that is incorporated in sulfur-containing biomolecules. SiR is composed of two subunits: the SiR flavoprotein (SiRFP, α) and the SiR hemoprotein (SiRHP, β). Historically, the model for SiR holoenzyme assembly is that eight SiRFP subunits (α_8_) combine with four SiRHP subunits (β_4_) to form a heterododecameric (α_8_β_4_) complex (referred to here as a dodecamer for simplicity) (Siegel *et al*., 1973; Siegel *et al*., 1974).

Each subunit has been well characterized but how they work as a dodecamer remains under investigation. SiRFP is homologous to cytochrome P_450_ reductase (CPR) and other similar diflavin reductases (Gruez *et al*., 2000; Hubbard *et al*., 2001; Iyanagi *et al*., 2012; Haque *et al*., 2014). The domain structure of this class of reductases consists of a variable N-terminal region followed by a “tether” that joins it to a flavodoxin-like domain (Fld domain) that is connected through a “linker” to a ferredoxin/NAD^+^ reductase domain (FNR domain) interrupted by a connection domain (Tavolieri *et al*., 2019) (Figure 1). Computational and structural analysis of SiRFP and the homologous CPR predict that the tether and linker are flexible (Gruez *et al*., 2000; Askenasy *et al*., 2018; Tavolieri *et al*., 2019; Rwere *et al*., 2024; Ghazi Esfahani *et al*., 2025). SiRHP is a monomeric metalloenzyme that contains a coupled siroheme/Fe_4_S_4_ cluster five iron center when it is isolated from its SiRFP partner (Crane *et al*., 1995).

**Figure 1:**
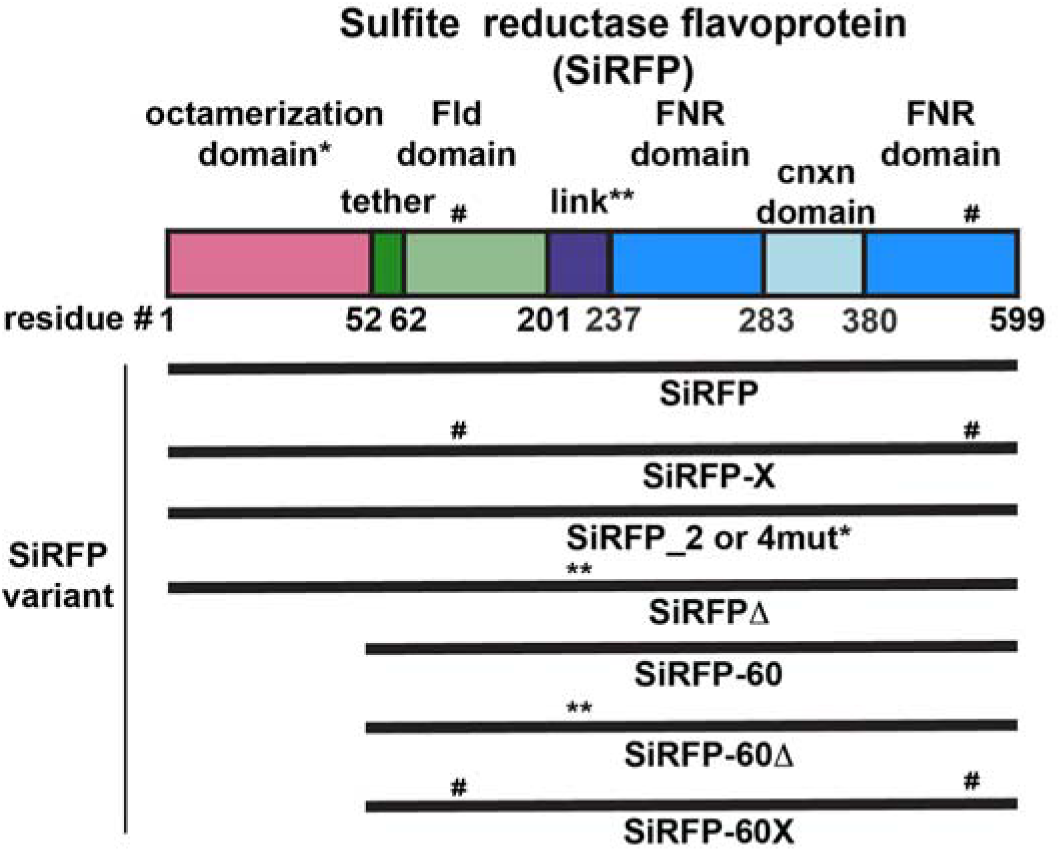
Domain architecture and residue alterations of well-characterized SiRFP variants. SiRFP is full length and octameric; SiRFP-X is a full length octamer with disulfide crosslinking; SiRFP_2 and SiRFP_4mut are the subject of this study; SiRFP-Δ is the octamer with a deletion in linker region; SiRFP-60 is monomer with deletion of the N-terminal 52 residues; SiRFP-60Δ is a monomer with deletion of the N-terminal 52 residues and in the linker region; SiRFP-60X is monomer with deletion of the N-terminal 52 residues and disulfide crosslinking. Deletions and disulfide crosslinking in specific domains are represented by *, ** and # respectively.

SiR uses three equivalents of NADPH to catalyze the six-electron reduction of SO_3_^2-^ to S^2-^. For each NADPH, two electrons sequentially reduce the flavin adenine dinucleotide (FAD) cofactor, then the flavin mononucleotide (FMN) cofactor, before moving to the to the iron center in SiRHP where the substrate binds at the distal face of the siroheme (Siegel *et al*., 1974). A well-characterized conformational change, mediated by the linker between the Fld and FNR domains, brings the flavin cofactors in proximity for electron transfer from the reduced hydroquinone FADH_2_ cofactor to the FMN (Freeman *et al*., 2017; Tavolieri *et al*., 2019; Murray *et al*., 2021; Murray *et al*., 2022; Walia *et al*., 2023; Ghazi Esfahani *et al*., 2025). The now reduced Fld domain repositions to deliver the electrons to the SiRHP active site for catalysis through an ill-defined mechanism. How these dynamic interactions between the SiRFP domains, and then with SiRHP, occur in the context of the α_8_β_4_ dodecamer remains something of a mystery because the region that connects the octamerization domain to the Fld domain has properties of intrinsic disorder that challenge traditional structure determination methods (Askenasy *et al*., 2018).

Despite the technical challenges for structural studies of flexible protein complexes, protein engineering combined with X-ray crystallography, small angle neutron scattering (SANS), proteolytic mass spectrometry, and cryo-EM analyses have revealed the following properties of the SiR complex (Figure 1 and Table 1). 1) Removing the first 51 residues of SiRFP result in a 60 kDa monomer (SiRFP-60) that binds in a 1:1 fashion with SiRHP (Zeghouf *et al*., 2000; Askenasy *et al*., 2018; Murray *et al*., 2021; Ghazi Esfahani *et al*., 2025). 2) An internal truncation of the linker joining SiRFP’s Fld and FNR domains (Δ-AAPSQS, residues 212-217) stabilizes the structural relationship between the domains in either the SiRFP monomer (SiRFP-60Δ (Tavolieri *et al*., 2019)) or octamer (SiRFP-Δ (Murray *et al*., 2022)). 3) Engineered disulfide crosslinks between the Fld and FNR domains rigidify SiRFP-60 (SiRFP-60X (Ghazi Esfahani *et al*., 2025)) but form both inter- and intra-molecularly through domain crossover in the context of the octamer (SiRFP-X (Walia *et al*., 2023)).

**Table 1:** Sulfite Reductase Flavoprotein (SiRFP) variants.

| Variant name | Variation | Property | Reference |
| --- | --- | --- | --- |
| SiRFP | Full-length, wild type | 536 kDa octamer | (Siegel and Davis, 1974) |
| SiRFP-60 | N-terminal 52 amino acid truncation | 60 kDa monomer | (Zeghouf *et al.*, 1998) |
| SiRFP-Δ | Internal deletion of residues 212-217 | 535 kDa octamer | (Tavolieri *et al.*, 2019) |
| SiRFP-60Δ | N-terminal 52 amino acid truncation/Internal deletion of residues 212-217 | 59 kDa monomer | (Murray *et al.*, 2021) |
| SiRFP-X | Full-length/C162T, C552S, W121C, N556C | Crosslinked, 536 kDa octamer | (Walia *et al.*, 2023) |
| SiRFP-60X | N-terminal 51 amino acid truncation/C162T, C552S, W121C, N556C | Crosslinked, 60 kDa monomer | (Ghazi Esfahani *et al.*, 2025) |
| NFPSh | SiRFP 1-52 – SUMO – 6xhis | 176 kDa octamer | This study |
| SiRFP_2mut | Y39A and F40A | mixed | This study |
| SiRFP_4mut | Q22A, Y39A, F40A, and Q47A | mixed | This study |

Small angle neutron scattering (SANS) studies on SiR, SiR-X (SiR with the disulfide crosslinked SiRFP octamer), SiRFP, and SiRFP-X further develop the model for SiR structure in which the SiRFP N-terminus is a central core, bringing together the N-terminal Fld domains at the center of a 536 kDa octamer (Walia *et al*., 2023). In the dodecamer, the four SiRHP subunits bind independently and at the periphery of the SiRFP octamer to generate an 800 kDa holoenzyme complex (Murray *et al*., 2022; Walia *et al*., 2023). The nature of the octamerization domain is unknown because the role of the N-terminus differs between diflavin reductase homologs. For example, eukaryotic microsomal CPR is a monomer that is tethered to the endoplasmic reticulum membrane through a 6 kDa N-terminal helical extension (Xia *et al*., 2019). In nitric oxide synthase, the diflavin reductase domain is expressed as a C-terminal fusion with a pterin binding oxygenase, sometimes with an intervening calmodulin binding regulatory domain (Campbell *et al*., 2014; Haque *et al*., 2014).

Here, we use an ensemble of complementary biophysical methods, including *tandem*-trapped ion mobility spectrometry/mass spectrometry (Tandem-TIMS), chromatography, structure modeling/mutagenesis, analytical ultracentrifugation (AUC), mass photometry, dynamic light scattering (DLS), and SANS, to analyze the role of the N-terminus in SiRFP assembly. We show that the SiRFP’s N-terminus is sufficient to mediate octamerization and identify four residues (Tyr39, Phe40, Gln22, and Gln47) that stabilize this assembly. Altering these side chains results in mixed-state variants of SiRFP with concentration-dependent oligomeric states. We further show that SiRFP’s N-terminus has ultra-stable properties even in the presence of strong denaturants and temperatures up to 70 °C. This stable higher order oligomerization property is rarely found in protein systems, suggesting this peptide could be used as a tool for studying other smaller protein systems.

## Results

### Full-length SiRFP forms an octamer

The asymmetric nature of the SiR dodecamer is curious because the 1:1 SiRFP:SiRHP minimal dimer remains active (Zeghouf *et al*., 2000; Tavolieri *et al*., 2019), leading some to propose that the α_8_β_4_ hypothesis is not accurate (Zeghouf *et al*., 2000). The highly flexible nature of SiRFP’s tether and linker complicates the interpretation of many biophysical techniques used to assess protein complex mass and stoichiometry that rely on the assumption that proteins are primarily globular, like size exclusion chromatography (SEC) (Hong *et al*., 2012), density sedimentation by AUC (Edwards *et al*., 2020), small angle scattering (SAXS or SANS) (Receveur-Brechot and Durand, 2012), or blue native polyacrylamide gel electrophoresis (BN-PAGE) (Gallagher, 2001).

To address the assumption that SiRFP spontaneously assembles to an octamer, the measurement of which could be influenced by its flexibility (Siegel and Davis, 1974), we used Tandem-TIMS (Liu *et al*., 2018) to directly measure the MW of full-length SiRFP that natively assembles into an octamer (Askenasy *et al*., 2015). The predominant species is 575.2 kDa, consistent with an octamer of full-length SiRFP (including the N-terminal six-histidine (six-his)-containing extension) that has eight FMN and eight FAD cofactors (Figure 2a and Table 2). The theoretical mass of the octamer is 575.3 kDa, a difference of ∼7 Da/protomer from the measured mass. A secondary species of 573.6 kDa (a difference of 155 Da/protomer) could arise from loss of the N-terminal-most methionine within the six-his tag (the molecular mass of methionine is 149 Da). C-terminal degradation is unlikely as the terminal tyrosine is involved in FAD binding (Tavolieri *et al*., 2019). UV/vis spectroscopy shows that recombinantly-expressed, N-terminal six-his SiRFP has a full complement of flavins (molecular mass of 785 Da for FAD and 456 Da for FMN) (Askenasy *et al*., 2015). SiRFP did not fragment into smaller subcomplexes in the gas phase.

**Figure 2:**
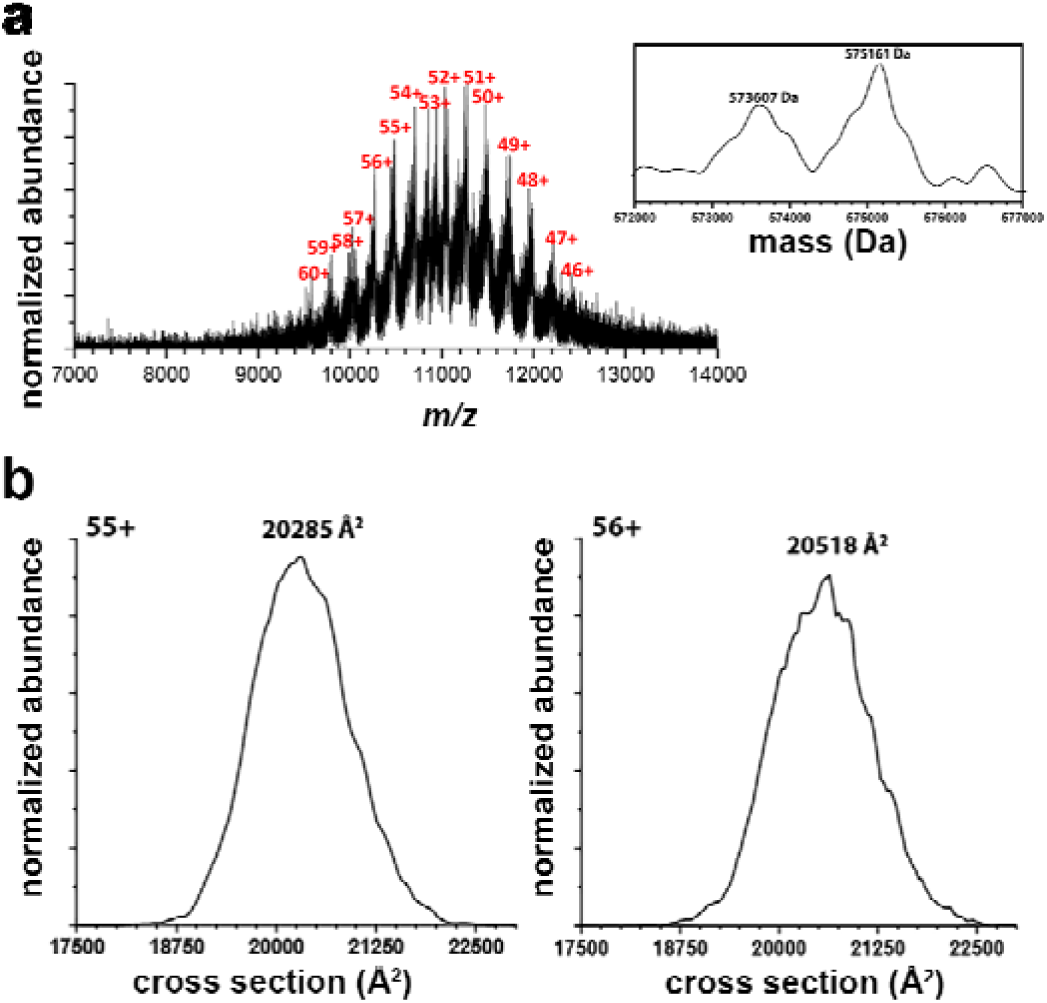
Tandem-TIMS/MS of 6xhis-SiRFP shows it is an octamer of 70.7 monomers in solution. **a.** Charge states (red) of the SiRFP octamer. **b.** Cross section (Å^2^) analysis of the two dominant charge states show the rotationally averaged area at the plane of the center of gravity.

**Table 2:**
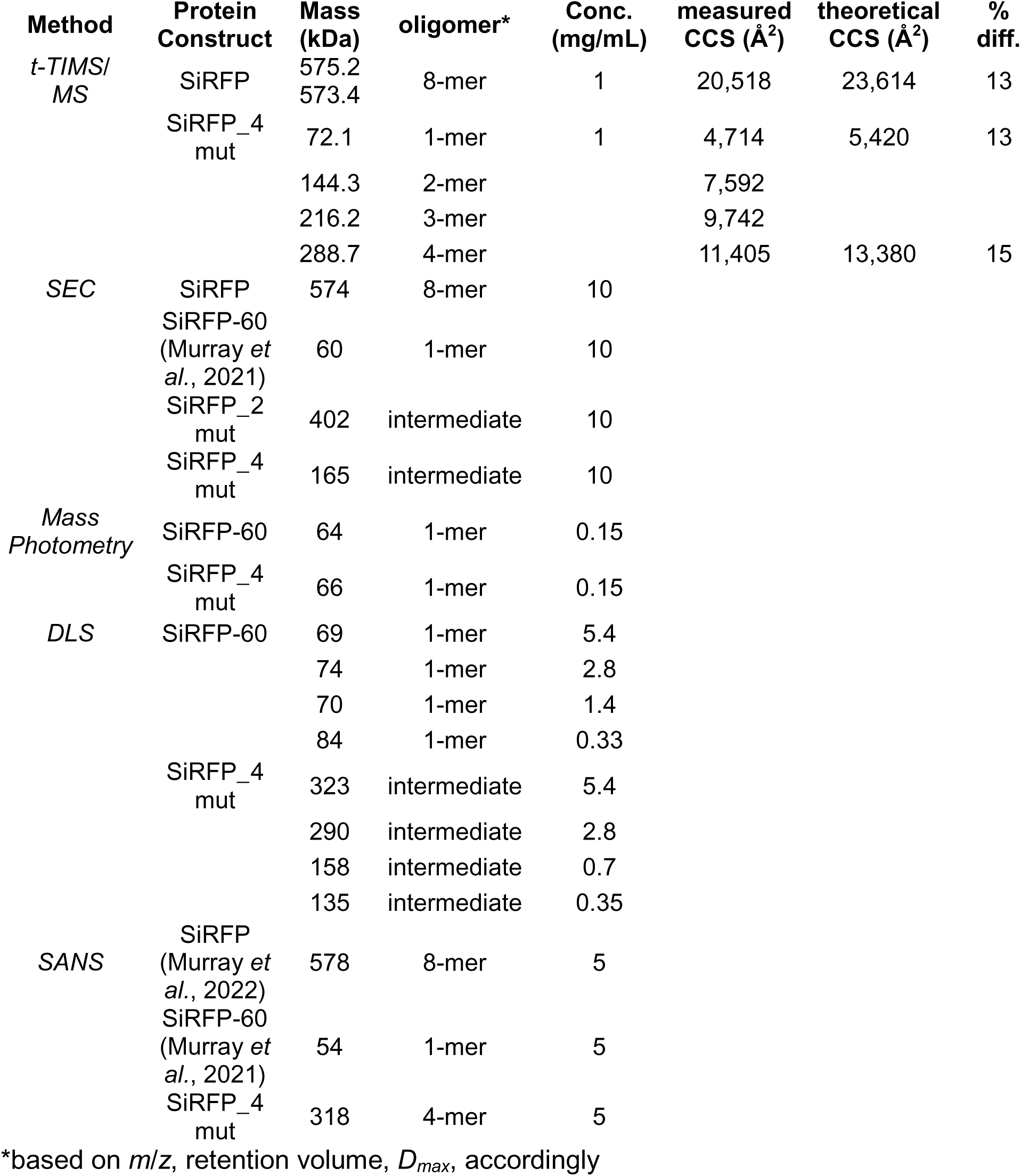
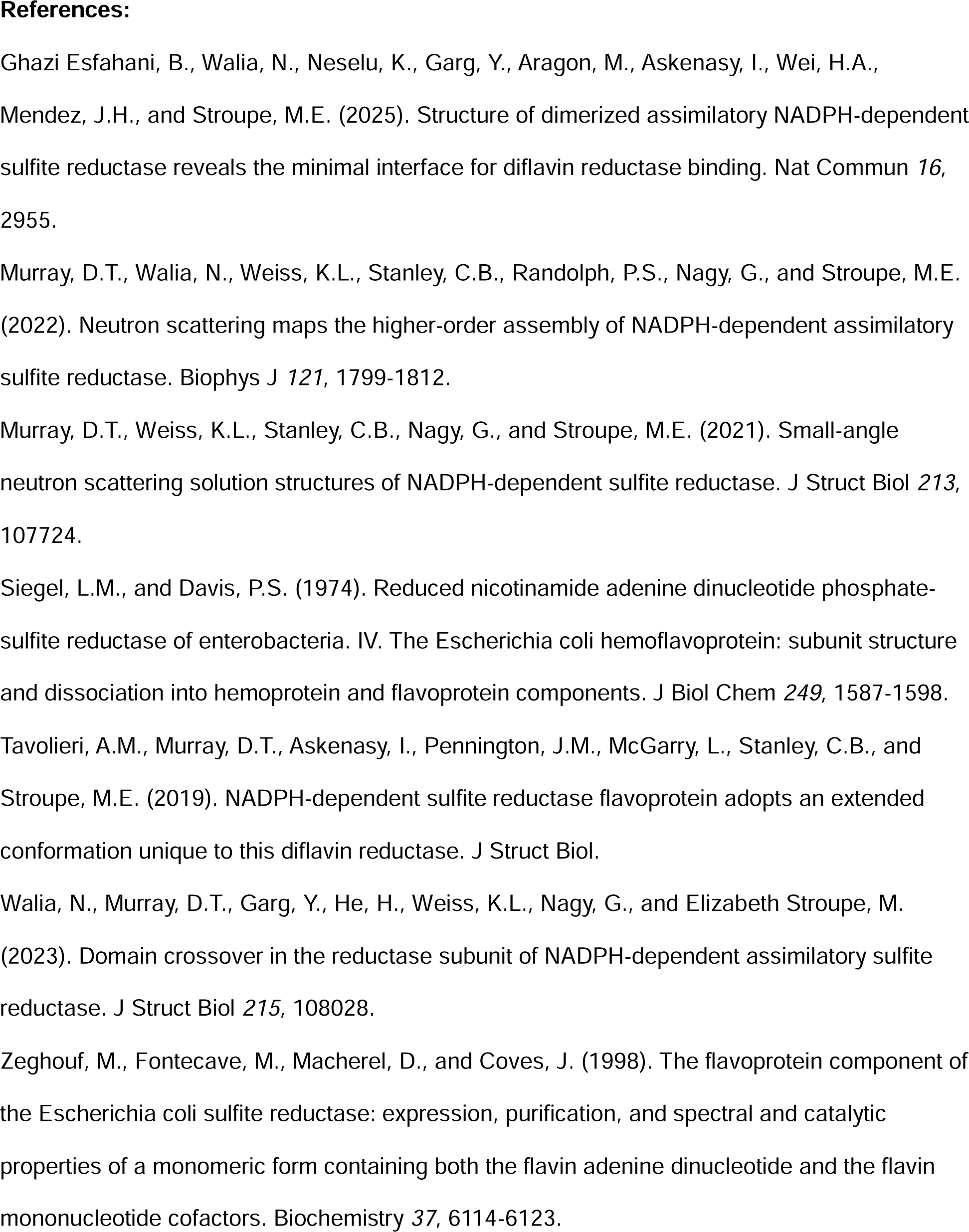
Summary of mass/oligomeric state determined by different techniques.

The ion mobility of a complex species is related to its collision cross section, thus shape information about the molecular structure of a species can be inferred from the Tandem-TIMS analysis (Liu *et al*., 2018). The measured collision cross section of the higher mass SiRFP octamer species, 20518 Å^2^, is consistent with theoretical rotationally averaged cross section calculated from a model based on prior SANS analysis of the same construct (Walia *et al*., 2023), 23614 ± 269 Å^2^ (Table 2).

### The N-terminus of SiRFP is sufficient for self-assembly into an octamer

SiRFP-60, the 60 kDa monomeric variant that lacks its first 51 residues, catalyzes NADPH-dependent reduction of the FAD and FMN cofactors that subsequently reduce SiRHP or a cytochrome *c* partner (Zeghouf *et al*., 1998; Askenasy *et al*., 2018). We do not know the role of those residues in holoenzyme assembly because the only structures of SiRFP are of SiRFP-60 variants lacking the N-terminus (Gruez *et al*., 2000; Tavolieri *et al*., 2019; Ghazi Esfahani *et al*., 2025). To test if the peptide is sufficient for octamerization, we constructed a chimera of SiRFP’s first 52 residues fused to a SUMO tag (NFPSh: <u>N</u>-terminus SiR<u>FP</u> with <u>S</u>UMO and <u>H</u>is tag) (Figure 3a). SUMO is typically used as a monomeric tag to increase the solubility of recombinantly expressed proteins (Kuo *et al*., 2014) but here we are using it as a chimeric carrier for the N-terminus of SiRFP.

**Figure 3:**
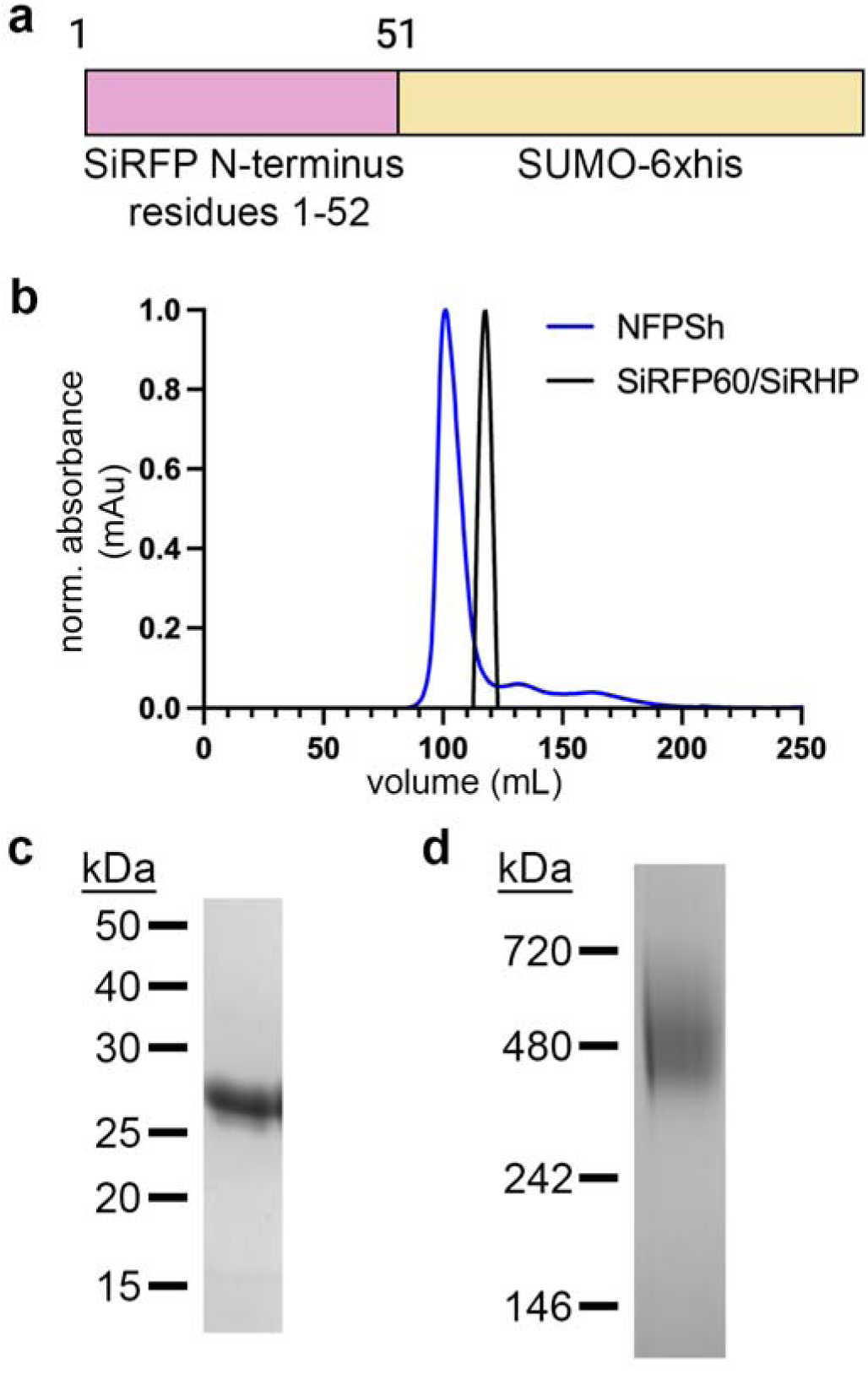
SiRFP’s N-terminal 52 residues are sufficient for octamerization. **a.** Schematic representing the NFPSh chimera. The SiRFP N-terminal region is pink and the SUMO moiety with a C-terminal 6xhis tag is yellow. **b.** The SEC elution profile of NFPSh (blue) shows an elution volume of 102 mL. As reference, the SiRFP/SiRHP heterodimer, which is ∼124 kDa in mass (Ghazi Esfahani *et al*., 2025), shows an elution volume of 120 mL (black), confirming that NFPSh does not behave as a monomer. **c.** SDS-PAGE gel analysis shows that, when denatured, NFPSh is ∼25 kDa (molecular weight (MW) markers in kDa). **d.** Blue Native gel analysis shows a higher MW of ∼450 kDa (MW markers in kDa).

Purified NFPSh fractionates earlier via SEC than the known 124 kDa SiRFP-60/SiRHP heterodimer (Ghazi Esfahani *et al*., 2025) (Figure 3b). Octameric NFPSh has a theoretical mass of 360 kDa, significantly larger than its theoretical monomeric weight of ∼27 kDa. SDS-PAGE analysis shows a denatured monomer close to 25 kDa in mass; however, BN-PAGE (Wittig *et al*., 2006) shows a higher molecular weight indicating that SiRFP’s N-terminus is sufficient for higher-order assembly of the octamer regardless of its C-terminal fusion (Figures 3c and d). Predicted disorder in the regions surrounding the components responsible for oligomerization likely accounts for the aberrantly high position of NFPSh on the native gel – the same phenomenon occurs for the wild type SiRFP octamer (Askenasy *et al*., 2015).

### The SiRFP N-terminal assembly is ultra stable

Given the lack of subcomplex fragmenting of SiRFP in the gas phase, we next explored the stability of the complex in solution using NFPSh as a model system to confine the analysis to the function of the octamerization domain. First, we tested our prediction that the N-terminal assembly was dominated by the hydrophobic stacking of Tyr39 and Phe40. We applied a mixture of NFPSh in 5% or 10% DMSO to a SEC column to determine if the presence of a hydrophobic solvent, DMSO, affected the oligomeric state of octameric SUMO. The elution volume of NFPSh was not significantly altered (Figure 4a).

**Figure 4:**
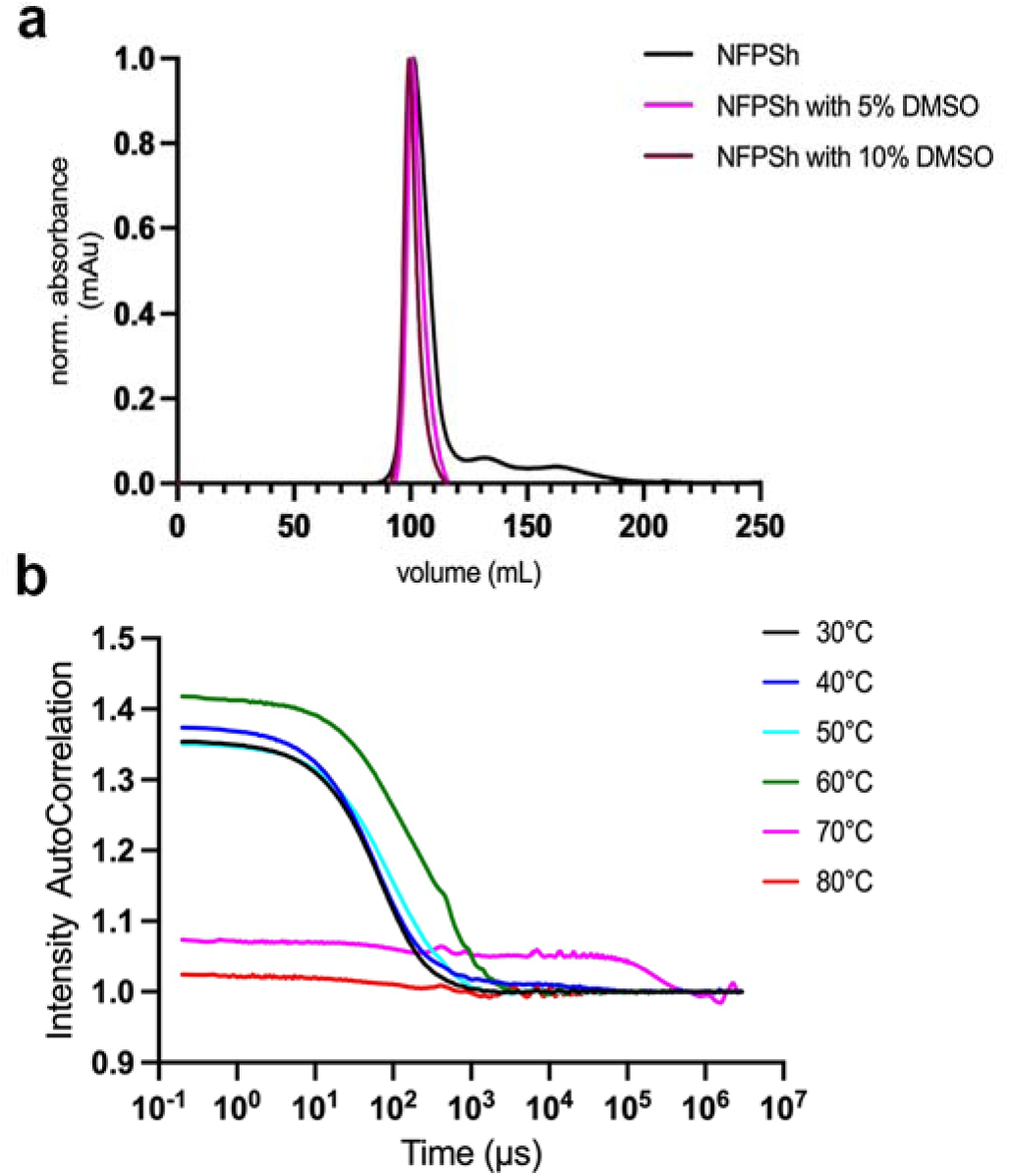
The N-terminus of SiRFP is ultra stable. **a.** DMSO has no effect on the oligomeric state of NFPSh. SEC profile of NFPSh with 5 or 10% DMSO shows no change in the elution volume suggesting no dissociation of the octameric helical bundle under mild hydrophobicity. **b.** DLS-measured thermal melting shows that NFPSh remains stable up to 70 °C.

We further tested the impact of other denaturants on the complex. We incubated NFPSh with a series of chaotropic agents (urea and guanidine hydrochloride (GdnHCl)), salt (KCl), and detergents (SDS, Tween-20, Triton-X, CHAPS, or Fos-choline 8 (FC8)) (Supplemental Figure 1). Finally, thermal denaturation of NFPSh, assessed with DLS, shows that the assembly remains intact up to 70 °C (Figure 4b). Together, these results indicate that the N-terminal helical bundle shows highly stable properties and remains intact even in the presence of strong denaturants.

### Critical residues in forming an octameric SiRFP

Given that we do not have a structure of SiRFP’s N-terminal peptide, we used AlphaFold (Jumper *et al*., 2021) to model the full-length structure of a single SiRFP polypeptide chain. AlphaFold predicts that the first 52 residues adopt short, disordered regions surrounding a short helix-turn-helix (Figure 5a).

**Figure 5:**
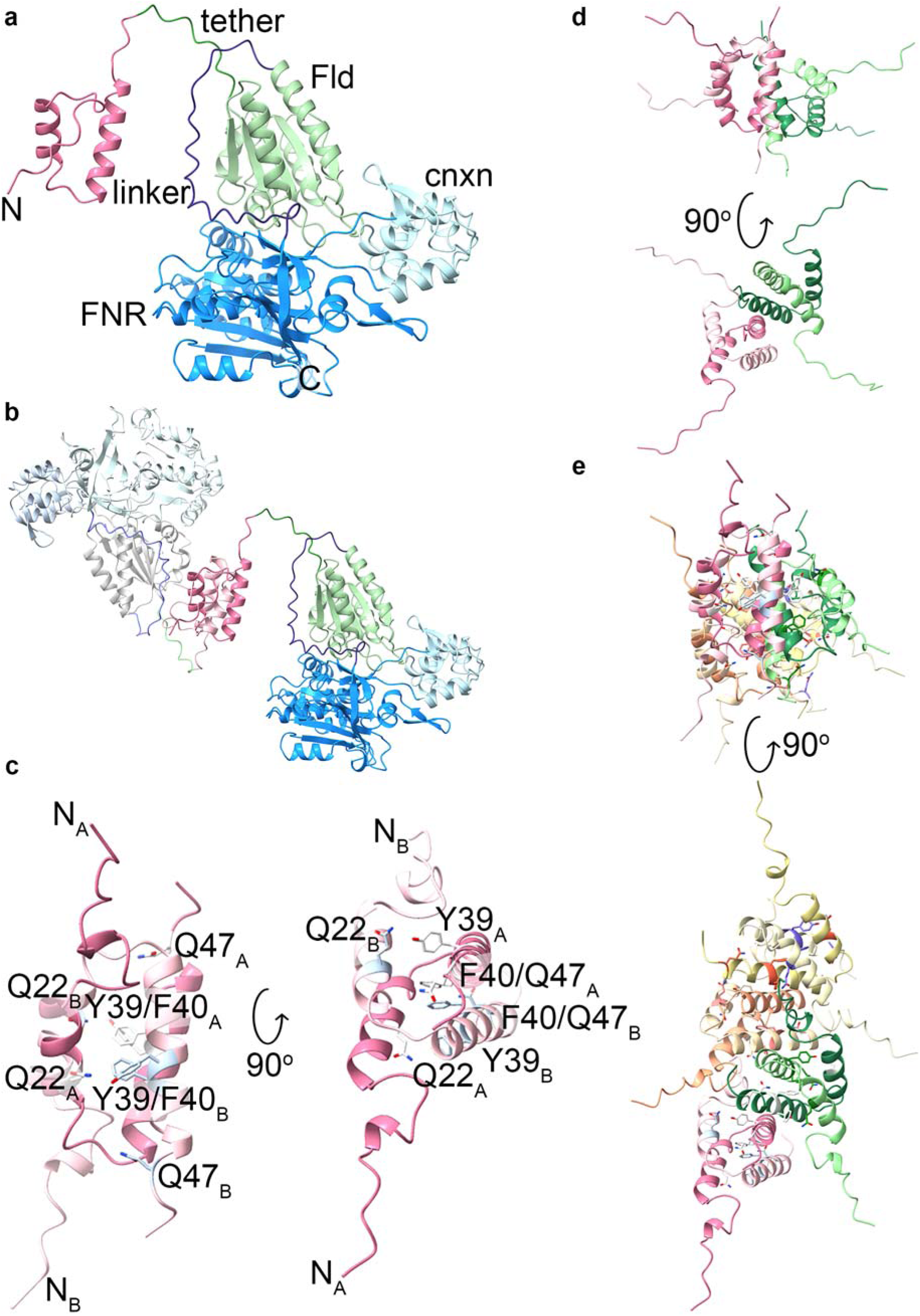
Modeling of SiRFP’s N-terminus and design of SiRFP variants. **a.** Molecular modeling of full-length SiRFP suggests the N-terminus consists of a short helix-turn-helix region (dark pink) followed by a tether (dark green), the folded Fld domain (light green), a 36 residue-long linker (purple), and then an FNR domain (dark blue) interrupted by a connection domain (cyan). Colors as in Figure 1. **b.** Two copies can assemble into a homodimer formed by a helical bundle. **c.** Four residues (Gln22, Tyr39, Phe40, and Gln47) are predicted as important in assembly. **d.** Dimers of helical bundles could form a tetramer. **e.** dimers of tetramers could form octamers.

We next aimed to understand how the predicted helix-turn-helix might oligomerize. With the AlphaFold-generated model of full-length SiRFP, we used a deep assembly algorithm implemented in HDOCK (Yan *et al*., 2020) to dock two subunits of SiRFP (Figure 5b). Two aromatic residues, Tyr39 and Phe40, were implicated as capable of π-π stacking with the symmetric partner from another subunit (Figure 5c). Two additional residues, Gln22 and Gln47 could make close contact with the symmetric residue from another subunit (Figures 5d and e). We thus hypothesized that altering either both Tyr39 and Phe40 or each of Gln22, Tyr39, Phe40 and Gln47 could result in full-length, but monomeric, SiRFP.

### SiRFP assembly variants are functionally active, but the octamer is disrupted

To test our hypothesis that these residues are responsible for the assembly of the SiRFP octamer, we generated two variants (Tables 1 and S1): one in which the aromatic side chains were altered to alanine (SiRFP_Y39A_F40A: SiRFP_2mut) and one in which all four side chains were altered to alanine (SiRFP_Y39A_F40A_Q22A_Q47A: SiRFP_4mut).

We used our established complementation test to assess if SiRFP_2mut or SiRFP_4mut remains functional. Both variants complement *cysJ*-deficient *E. coli* (Baba *et al*., 2006) when grown under limiting sulfur conditions (Figure 6a). Further, both SiRFP variants are soluble when purified with an established expression/purification protocol (Askenasy *et al*., 2018). UV-vis spectroscopy shows the purified variants have signals nearly identical to the wild type enzyme, indicating that they remain properly folded to bind the flavin cofactors (Supplemental Figure 2).

**Figure 6:**
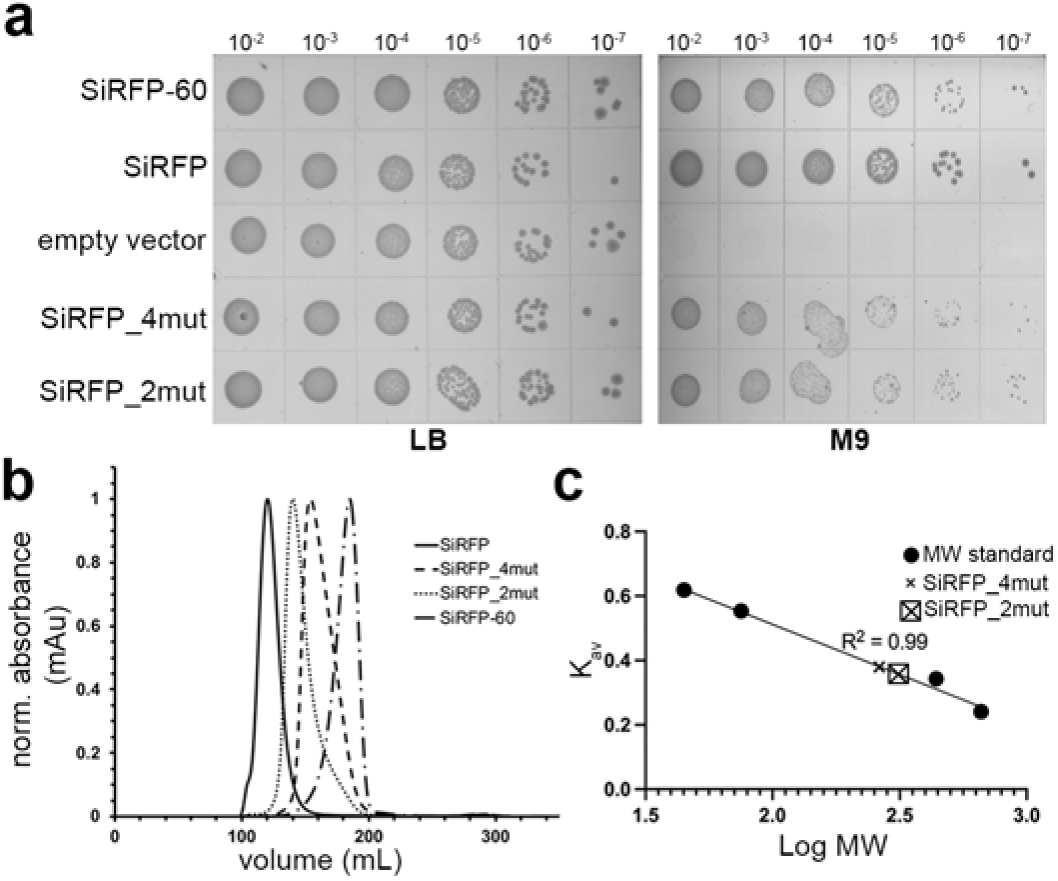
SiRFP function is not impacted by N-terminal variants but the same variations impact oligomeric state. **a.** SiRFP_4mut and SiRFP_2mut complement SiRFP-deficient *E. coli* in the absence of reduced sulfur at the same levels of wild-type SiRFP and monomeric SiRFP-60. **b.** SEC analysis shows that SiRFP_4mut and SiRFP_2mut are intermediate in mass between octameric SiRFP and monomeric SiRFP-60 but run as broad species. **c.** Molecular weight standards to calibrate the SEC column.

The SEC profiles confirm that neither variant behaves as either an octamer or monomer when compared to full-length WT SiRFP or monomeric SiRFP-60 (Figure 6b and Table 2). Specifically, each variant elutes as a broad peak, suggesting an intermediate oligomerization state that is higher than a monomer but not a full octamer. By SEC, the average SiRFP_2mut species eluted as an oligomer of 402 kDa, consistent with an oligomer of about 5.6 protomers (Figure 6c). The average SiRFP_4mut species eluted as an oligomer of 263 kDa, consistent with an oligomer of about 3.6 protomers (Figure 6c).

### The disrupted oligomeric state is concentration dependent

BN-PAGE supports the conclusion that the amino acid variations disrupt SiRFP assembly (Supplemental Figure 3). SiRFP_2mut assembles as a heterogenous species, which explains its broad SEC peak. The SiRFP_4mut variant predominantly showed a monomeric state. All further analysis focused on SiRFP_4mut.

We next performed AUC and mass photometry to clarify the stoichiometry of SiRFP_4mut, using SiRFP-60 as a marker for monomeric SiRFP (Supplemental Figure 4 and Table 2). Each variant sedimented with an envelope consistent with a predominantly monomeric state: an S value of 3.8, as expected from the previous analysis (Murray *et al*., 2021; Murray *et al*., 2022). SiRFP-60, well characterized as a monomer (Gruez *et al*., 2000), showed 96% of the input as a monomer of 68 kDa. SiRFP_4mut showed a predominant species at 69 kDa (89%), but with varied higher-order assemblies as minor species (dimer: 136 kDa, 5.6%; trimer: 232 kDa, 2%, and tetramer: 340 kDa, 1%). Mass photometry supported the interpretation from AUC that SiRFP_4mut exists predominantly as a monomer: 97% of the SiRFP_4mut species were attributed to a 66 kDa species whereas for SiRFP-60, 99% were attributed to a 64 kDa species (Supplemental Figure 5).

DLS measurements were used to evaluate the oligomeric state and solution homogeneity of SiRFP-60 and SiRFP_4mut across a range of concentrations from 0.3 to 5.4 mg/mL (Supplemental Figure 6 and Table 2). For SiRFP-60, the hydrodynamic radius was narrowly distributed from 3.6 to 4 nm (mean 3.7 ± 0.1 nm) regardless of concentration. The corresponding apparent MW was estimated to be 68 to 83 kDa, consistent with a monomer. The polydispersity index (PDI) was 0.1 to 0.2, suggesting a monodisperse sample. In contrast, SiRFP_4mut showed concentration-dependence with a hydrodynamic radius of 4.9 to 7.0 nm (mean 5.9 ± 1.2 nm), corresponding to an apparent MW from 136–323 kDa. The PDI ranged from 0.2–0.3.

### SANS measurement shows that SiRFP_4mut forms a flexible, asymmetric tetramer that dominates its scattering profile

We next measured SANS on SiRFP_4mut. The scattering profiles are consistent with a monodisperse complex (Figure 7a) with an *R_g_* of 67.1 ± 0.5 Å and *D_max_* of 209 Å (Figure 7b). These measurements are smaller than octameric SiRFP (*R_g_* of 80.8 ± 0.8 Å and *D*_max_ of 260 Å, (Murray *et al*., 2022)), but almost double the *R_g_* and *D_max_* of SiRFP-60 (*R_g_*of 32.2 ± 0.1 Å and *D_max_*of 113 Å, (Murray *et al*., 2021)), suggesting that in solution and higher concentration SiRFP_4mut oligomerizes (Table 2). The Kratky plot shows similar features to that of octameric SiRFP (Murray *et al*., 2022), which is known to be particularly flexible (Supplemental Figure 7). Modeling of the scattering envelope by DENSS showed the possible arrangement of four subunits of SiRFP_4mut based on the HDOCK tetramer, suggesting an asymmetric arrangement of the subunits with a helical bundle at its core (Figure 7c).

**Figure 7:**
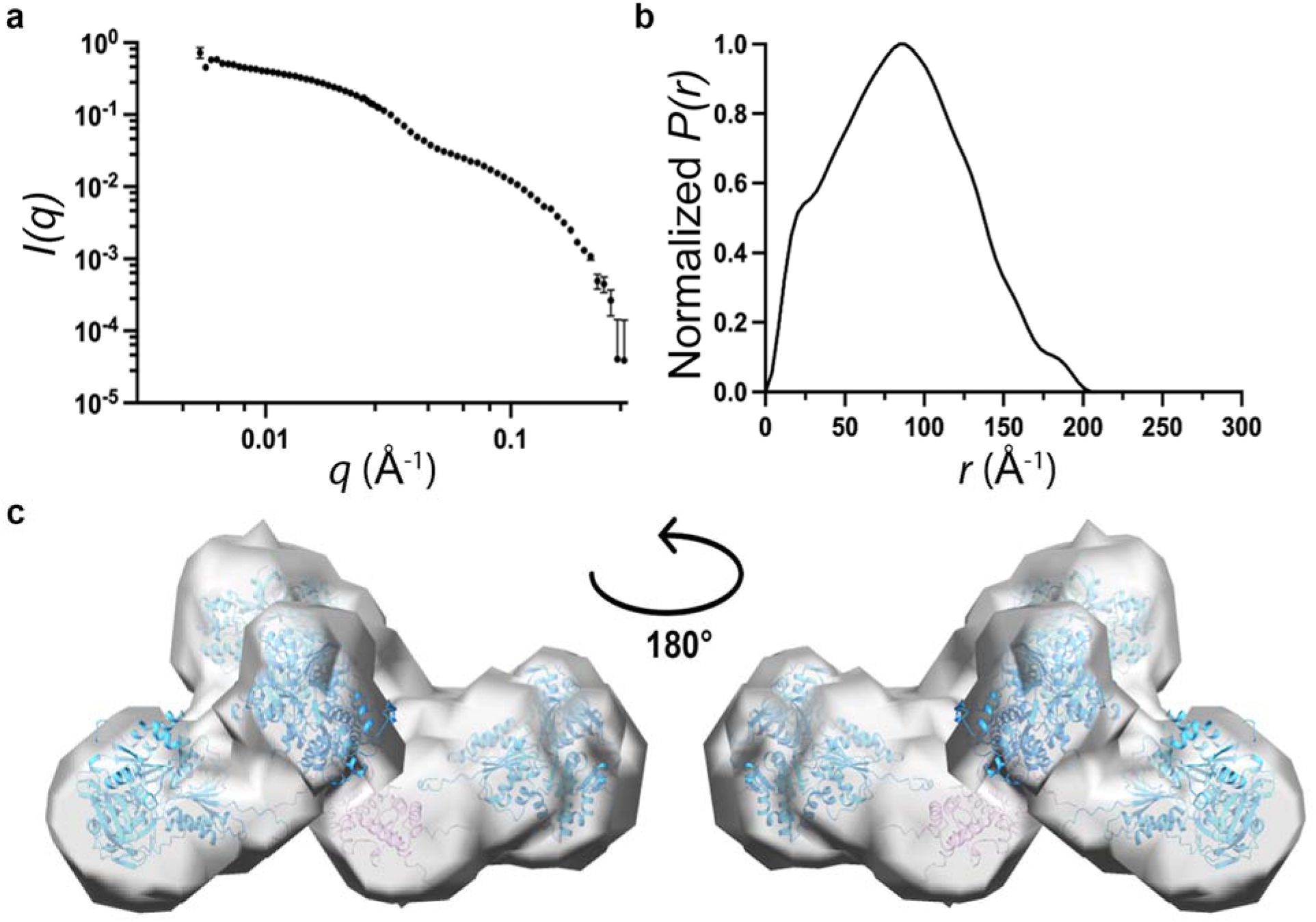
SANS measurements on SiRFP_4mut. **a.** Scattering profile of SiRFP_4mut. **b.** Distance distribution plot shows an *R_g_*of 67.1 ± 0.5 Å and *D_max_*of 209 Å. **c.** DENSS envelope of SiRFP_4mut and one possible arrangement of four subunits of SiRFP with the N-terminal helical bundle.

### Tandem-TIMS on SiRFP_4mut suggest a mixture of oligomeric states in solution

To reconcile the mixed results described above, we used Tandem-TIMS to measure the behavior of SiRFP_4mut using a method that is less biased by the behavior of flexible protein complexes in solution. The predominant species behaves as a monomer, consistent with AUC, mass photometry, and DLS at lower concentrations. A measurable component that corresponds to dimer, trimer, and tetramer was also observed by Tandem-TIMS, reconciling the DLS measurements at higher concentration, SEC, and SANS (Figure 8a and b and Table 2). The measured collision cross-section of each species shows increasing area as SiRFP_4mut’s predicted oligomeric states increase (Figure 8c and Table 2). The cross section of the predominant monomeric species (4714/4720 Å^2^) compares with the theoretical cross section calculated from the X-ray crystal structure of SiRFP-60 (PDB ID 6VEF, (Tavolieri *et al*., 2019)), 5420 ± 37 Å^2^). Similarly, the cross section of the tetrameric species is consistent with the model based on the envelope derived from the SANS measurement (11,405 Å^2^ compared with 13,380 Å^2^). The 13-15% overestimation of the theoretical collisional cross section is typical for molecules with conformational flexibility (Bleiholder *et al*., 2011).

**Figure 8:**
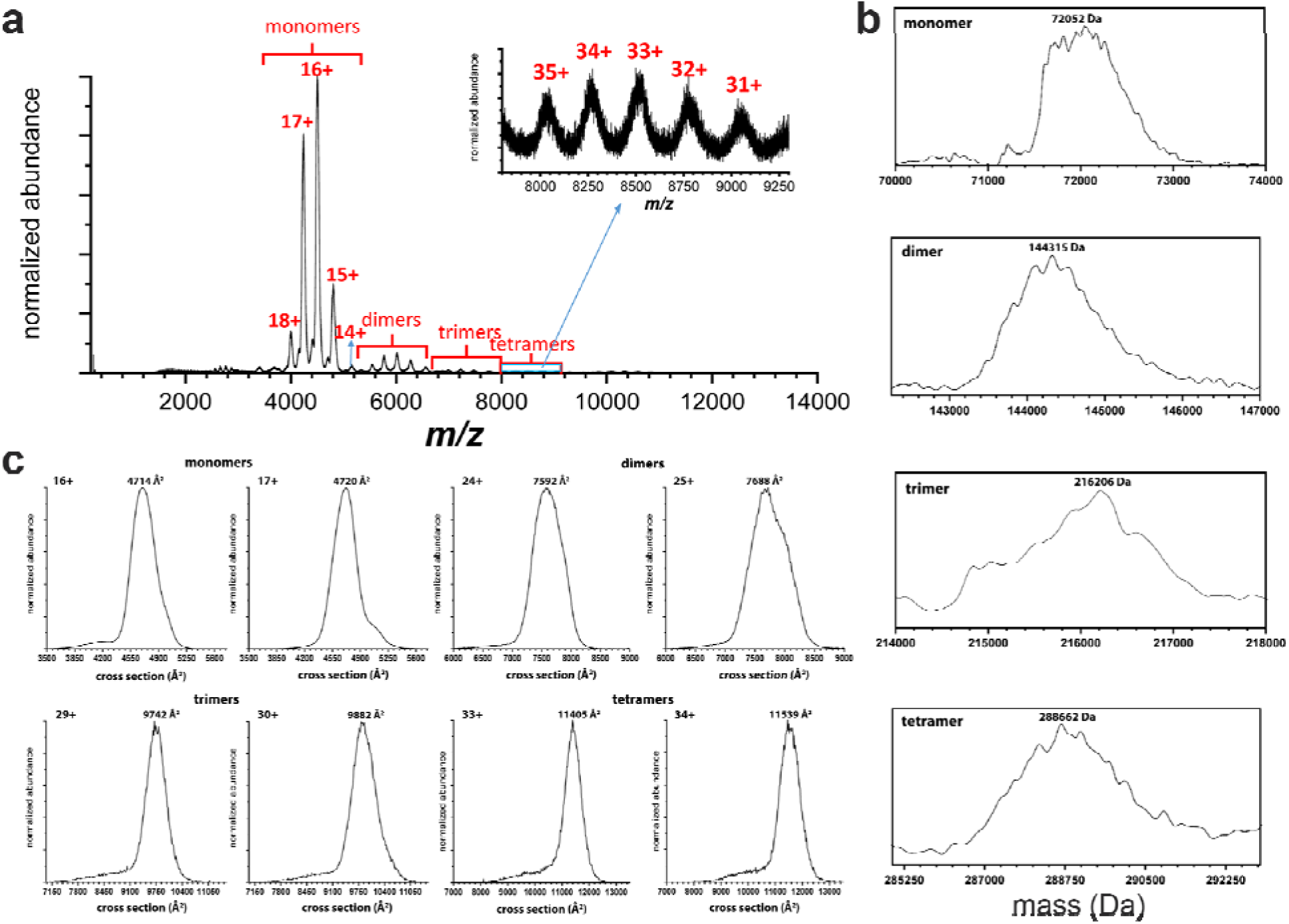
Tandem-TIMS/MS on SiRFP_4mut. **a.** There is a mixture of oligomeric states of SiRFP_4mut. **b.** Mass determination of each of the four oligomeric states. **c.** Cross section analysis of the SiRFP_4mut states.

## Discussion

SiR’s hetero-dodecameric state (α_8_β_4_), where 8 copies of SiRFP assemble with 4 copies of SiRHP, was first predicted upon SiR’s isolation in the 1970s (Siegel *et al*., 1973) but remains a curiosity because a minimal dimer of SiRFP and SiRHP retains activity (Zeghouf *et al*., 2000; Tavolieri *et al*., 2019; Ghazi Esfahani *et al*., 2025). Experiments aimed to disprove the α_8_β_4_ hypothesis have not provided evidence for an alternative model (Askenasy *et al*., 2015; Murray *et al*., 2022; Walia *et al*., 2023). Nevertheless, high-resolution structure determination of the SiRFP octamer has been unsuccessful thus far owing to its highly flexible nature. Together, these observations define an unanswered question that motivated this study: what is the nature of SiRFP’s spontaneous assembly into such a stable octamer?

Confirming the octameric nature of SiRFP was our first objective. Defining protein oligomeric state by biophysical methods like native gel electrophoresis, solution scattering, or SEC analysis, can be misleading if the molecule is highly flexible or the complex formation is concentration dependent. Tandem-TIMS largely avoids complications from flexibility because the mass of intact protein complexes is analyzed in the gas phase. In this way, we show that the SiRFP oligomer is octameric.

Tandem-TIMS provides another advantage in studying protein complexes: secondary fragmentation of protein oligomers by ion fragmentation can provide clues into the stable assemblies that come together to form the native complex. By this analysis, we identified another curious property of the native SiRFP octamer: it does not fragment into smaller assembly units like tetramers or dimers. In fact, it does not even fragment into monomers.

We show for the first time that SiRFP’s N-terminal 1-51 residues are sufficient, not just necessary, for oligomerization into an octamer because fusion of the peptide to the unrelated SUMO protein is sufficient to convert SUMO from a monomer to an octamer. This observation supports the understanding that the canonical diflavin reductase Fld-linker-FNR/connection domain architecture folds independently from the N-terminal peptide, which is idiosyncratic to other diflavin reductase family members. Curiously, assembly through these 52 residues is resistant to disassembly under a series of denaturing agents, including chaotropic agents, salts, detergents, or temperature.

This observation led us to model the peptide region to identify residues that are possibly responsible for its self-assembly. We identified four residues, Tyr39, Gln22, Phe40, and Gln47, as candidates for mediating oligomerization. Specifically, altering those residues to alanine breaks the octamer, forming a mixed species of full-length SiRFP monomers, dimers, trimers, and tetramers that retain the ability to reduce SiRHP to form reduced sulfur and support bacterial growth in the absence of supplied S^2-^. The mixed oligomeric state of the SiRFP_4mut variant suggests that weakened subunit-subunit interactions results in a dynamic equilibrium between the subunits, unlike in the stable wild type octamer. Perhaps this is one underlying feature of the asymmetric dodecamer that facilitates the efficient six electron reduction of sulfite for sulfur assimilation.

SANS measurements of SiRFP_4mut are consistent with an assembly that is intermediate between monomeric and wild type octameric SiRFP. Altering all four of the residues we identified from modeling the octamer (Tyr39, Gln22, Phe40, and Gln47) is necessary to “break” the octamer – only altering the hydrophobic residues, as in SiRFP_2mut, resulted in a partially dissociated oligomer. The genetic structure of the *cys* operon in *E. coli*, where the genes encoding SiRFP (encoded by *cysJ*, the α subunit) and SiRHP (encoded by *cysI*, the β subunit) are only present in a single copy, raises the question of how the asymmetric α_8_β_4_ dodecamer assembles. The observation that SiRFP_2mut partially dissociates suggests that assembly of the octamer occurs in a stepwise fashion as the ribosome translates multiple copies of the *cysJ* mRNA.

Together, these properties make SiRFP’s N-terminal domain a suitable tag for fusing with smaller molecular weight proteins to oligomerize the smaller proteins into an octamer, perhaps enhancing them as potential candidates for structural studies that rely on larger complexes like cryo-EM.

## Materials and Methods

### Generation of SiRFP variants

Codons for residues 53-599 were truncated from the *cysJ* gene, leaving behind only codons for residues 1-52, which encode the SiRFP octamerization domain. The sequence encoding SUMO was added C-terminally, followed by codons for a six-histidine tag, and the chimera was cloned into a pET14b vector (Millipore Sigma, Burlington, MA, USA). We call this chimera “N-terminus Flavoprotein with SUMO and His tag (NFPSh)”. The rationale for fusing the SiRFP N-terminus with the SUMO tag is to enhance the solubility of the N-terminal peptide and to test its ability to oligomerize an unrelated protein.

N-terminal 6xhistidine-tagged *cysJ* encoded in the pBAD plasmid (Invitrogen, Waltham, MA, USA) is an established expression vector and was used as the template for generating two (Y39A and F40A, SiRFP_2mut) or four alterations (Q22A, Y39A, F40A, and Q47A, SiRFP_4mut) in the N-terminal region of the full-length SiRFP via Q5 mutagenesis (Tables 1 and S1).

### Protein expression and purification

All SiRFP variants were purified as previously described (Askenasy *et al*., 2015; Askenasy *et al*., 2018; Murray *et al*., 2022; Walia *et al*., 2023). SiRFP_2mut or SiRFP_4mut were generated and expressed from a pBAD vector (ThermoFisher Scientific, Waltham, MA, USA) (Table 1). NFPSh/pET14b was transformed into BL21(DE3) cells (ThermoFisher Scientific, Waltham, MA, USA) and SiRFP_2mut or SiRFP_4mut/pBAD were transformed into LMG194 cells (Invitrogen, Carlsbad, CA, USA). Either cell line was grown in LB broth at 37 °C in the presence of 100 μg/mL ampicillin. NFPSh-expressing cells were induced with 1 mM β-D-1-thiogalactopyranoside (IPTG) at 30 °C for 4 h and SiRFP_2mut or SiRFP_4mut-expressing cells were induced with 0.05% arabinose at 25 °C for 4 h. Each variant was purified using a combination of Ni-NTA affinity chromatography (Cytiva, Marlborough, MA, USA), anion exchange chromatography (Cytiva, Marlborough, MA, USA), and Sephacryl S200-HR SEC (Cytiva, Marlborough, MA, USA). Purified samples were polished by SEC in 50 mM potassium phosphate, pH 7.8, 100 mM NaCl, 1 mM EDTA.

### Tandem-TIMS

#### Sample preparation

Ammonium acetate (≥99.99%) and LC-MS grade water were obtained from Sigma-Aldrich (St. Louis, MO). SiRFP samples were desalted 1x at 14,000 x g for 5 min using centrifugal filter devices (30 kDa Amicon Ultra; Millipore Corporation, Saint. Louis, MO). ESI tuning mix was obtained for ion mobility calibration from Agilent (Santa Clara, CA).

#### Tandem-TIMS measurements

Ion mobility measurements were performed on a Tandem-TIMS-QqTOF (Liu *et al*., 2018; Kirk *et al*., 2019) (Figure S1) with nitrogen buffer gas. An aliquot of 0.2 µM of each sample prepared in 50 mM ammonium acetate was sprayed out of in-house fused silica pulled tips (∼1 µm tip) prepared using a laser-based micropipette puller (Sutter Instruments, Novato, CA) for electrospray ionization (ESI). Ions produced from ESI were mobility-separated in the first TIMS analyzer (TIMS-1), selected and collisionally activated in the interface region, and finally mobility-separated in TIMS-2 prior to mass analysis (Liu *et al*., 2018; Kirk *et al*., 2019). Cross sections were calibrated as described (Liu *et al*., 2018) using Agilent ESI tuning mix ions (*m/z* 922, 1522, 2122).

#### Tandem-TIMS settings

Buffer gas was infused into the instrument via the desolvation gas unit of the nano electrospray source with a flow rate of 2.5 L min^-1^. The ion accumulation time for the TIMS1 cartridge was set to 29.83 ms. The desolvation gas temperature was kept at 323 K and the electrospray voltage was set to 875 V. The pressures at p1/p2/p3/p4 (Figure S2) were kept at 3.1/1.8/0.9/0.4 mbar. RF potentials with a frequency of 566 kHz, 443 kHz, and 3.2 MHz are applied to the TIMS-1 analyzer, TIMS-2 analyzer, and the hexapole, respectively. The RF peak-to-peak amplitude was set to 250 V for both TIMS cartridges. All spectra were acquired in positive ion mode.

### Size Exclusion Chromatography

A Gel Filtration High Molecular Weight Calibration kit (Cytiva, Marlborough, MA, USA) was used according to the manufacturer’s protocol to standardize a Superose 6 10/300 column (Cytiva, Marlborough, MA, USA) in 50 mM MES, pH 6.9 buffer after running blue dextran to identify the void volume. SiRFP_4mut, injected in the same volume as the standards, was passed over the same column at the same flow rate and in the same buffer. The known MW standards were as the elution volume normalized for the void (K_av_) on the X-axis and the log of molecular weight on the Y-axis in Microsoft Excel, a linear equation was calculated and then used to determine the molecular weight of the SiRFP_4mut based on its K_av_.

### Dynamic Light Scattering Experiments

DLS measurements were performed on a DynaPro 99 instrument along with a DynaPro-MSXTC temperature-controlled microsampler (Wyatt Technology, Santa Barbara, CA, USA) to evaluate the oligomeric state and solution homogeneity of monomeric SiRFP-60 and SiRFP_4mut across concentrations (0.3 to 5.6 mg/mL). Stock protein (5.4 or 5.6 mg/mL for SiRFP-60 or SiRFP_4mut) was diluted into 50 mM potassium phosphate, pH 6 and centrifuged at 14000 x g for 10 min at 4 °C. NFPSh in the same buffer was heated gradually from 30-70 °C. Measurements were made in triplicate and analyzed using Dynamic version 7.0.0.94 software.

The Stokes-Einstein’s equation was used to calculate the hydrodynamic radius (R_H_) and the polydispersity percentage from the diffusion coefficient (D_t_).

R_H_ = kT/6πηD_t_

Where k represents the Boltzmann constant, T is temperature (in Kelvin) and η is the viscosity of water.

### Complementation survival assay

A complementation survival assay was performed on SiRFP_2mut or SiRFP_4mut as previously described (Askenasy *et al*., 2015; Askenasy *et al*., 2018; Walia *et al*., 2023). SiRFP deficient *E. coli cysJ*^−^ (Baba *et al*., 2006) were transformed separately with pBAD vectors encoding wild type, octameric SiRFP, monomeric SiRFP-60, empty pBAD vector, SiRFP_4mut, or SiRFP_2mut. Cells were grown overnight at 37 °C in the presence of 100 μg/mL ampicillin to select for the exogenous plasmid and 50 μg/mL kanamycin to select for the *cysJ*^−^ deficiency. Cells were harvested by centrifugation for 10 mins at 4,000 x g and washed with M9 salts. Cell densities were normalized based on the optical density measured at 600 nm, serial dilutions were prepared, and cells were plated on LB or M9 plates.

### Sedimentation Velocity AUC

The sedimentation velocity experiment was performed using a Beckman ProteomeLab XL-I Analytical Ultracentrifuge with An60-Ti rotor (Beckman Coulter, Brea, CA, USA) at Oak Ridge National Laboratory (ORNL). The samples were diluted to 0.5 mg/ml in 50 mM potassium phosphate, pH 7.8, 100 mM NaCl, and 1 mM EDTA buffer. The cells were assembled using sapphire windows and a rotation speed of 32,500 rpm was used to measure absorbance at 280 nm at 20 °C for 7 h. The data were analyzed in the Sedfit software (Schuck *et al*., 2014) and S values of the samples were calculated accordingly.

### Mass photometry measurements

Mass photometry experiments were performed on a TwoMP mass photometer (Refeyn Ltd., Oxford, England). Instrument focus was established with 18 μL of buffer (50 mM potassium phosphate, pH 6) before the addition and mixing of 2 μL of protein sample for data collection. Movies were recorded for 60 s with AcquireMP acquisition software. Bovine serum albumin (66, 132 kDa), lactate dehydrogenase (140 kDa), and thyroglobulin (660, 1320 kDa) were used as mass calibrants to generate a standard curve prior to measuring SiRFP-60 and SiRFP_4mut at 20DnM and 5 nM, respectively. Data analysis was performed using DiscoverMP software. Gaussian distributions were fit to the peaks to determine the average molecular mass, standard deviation, and skewness.

### SANS measurements

SANS measurements were performed on NFPSh and SiRFP_4mut as previously performed on other SiRFP variants (Murray *et al*., 2021; Murray *et al*., 2022; Walia *et al*., 2023). Each sample was dialyzed against 100% D_2_O-containing buffer (50 mM potassium phosphate, pH 7.8, 100 mM NaCl, 1 mM EDTA). Neutron scattering was collected on an Extended Q-Range Small-angle Neutron Scattering diffractometer (EQ-SANS, Beam Line-6) at the Spallation Neutron Source (SNS) at ORNL (Heller *et al*., 2018). Two configuration settings were used, as before, covering a sufficient q range (0.005-0.7 Å^-1^) 4 m sample-to-detector distance with 10-13.6 Å wavelength band and 2.5 m sample-to-detector distance with 2.5-6.4 Å wavelength band. Data were corrected for detector sensitivity, followed by subtraction of the empty cell and buffer scattering. Circular averaging was performed to generate one-dimensional scattering profiles using standard *drtsans* software (Heller *et al*., 2022) and then placed on an absolute scale using a calibrated porous silica standard.

### SANS data analysis and modeling

The BioXTAS RAW package (Hopkins *et al*., 2017) was used for SANS data analysis, including Guinier and dimensionless Kratky plot analysis. The real space *I(O)*, *R_g_* and *D_max_*were calculated using pairwise distance distribution *(P(r))* plots using GNOM, implemented in the ATSAS suite (Franke *et al*., 2017). *Ab initio* SANS models were calculated in DENSS operating in “slow” mode (Sumner and Qian, 2022). 20 reconstructions were generated and then averaged, aligned and refined without symmetry constraints. Data has been deposited in the Small Angle Scattering Biological Data Bank under the code SAS7590.

### Blue Native Gel Analysis

BN-PAGE analysis was performed as previously described for SiR. 2 μg of protein was mixed with the following denaturants: 0.5 or 1 M urea or GdnHCl, 2 M KCl, or 1-2% detergents (SDS, tween-20, triton-X, CHAPS, or FC8) and incubated on ice for 1 h. 4-16% NativePAGE gels were loaded with the sample/denaturant and a G-250 additive or the NativeMark size standard (ThermoFisher Scientific, Waltham, MA, USA). Standard anode and cathode buffers were used according to the manufacturer’s protocol for electrophoresis at 4 °C. Gels were run at 150 V for 1 h, then the voltage was increased to 250 V for 45 m until the current dropped below 4 mA. Gels were destained overnight in 40% methanol and 10% acetic acid to fix the proteins.

## Acknowledgements

We thank Dr.s Christopher Stroupe and Ratkim Roy for helpful discussions. A portion of this research used resources at Spallation Neutron Source, a DOE office of Science User Facility operated by the Oak Ridge National Laboratory. The beam time was allocated to EQ-SANS on proposal number IPTS-30534.The Office of Biological and Environmental Research also supported work at the ORNL Center for Structural Molecular Biology. This research was supported in part by an appointment to the Oak Ridge National Laboratory GRO Program, sponsored by the U.S. Department of Energy and administered by the Oak Ridge Institute for Science and Education. Further support came from National Science Foundation grants MCB1856502 and CHE1904612 to M.E.S. and CHE 2305173 to C.B. and F.C.L. and National Institutes of Health grant R01GM135682 to C. B..

