## Supplemental Figures for "Critical amino acid residues in the N-terminal domain of NADPH-dependent assimilatory sulfite reductase flavoprotein mediate octameric assembly"

91 Chieftain Way

Tallahassee, FL 32306

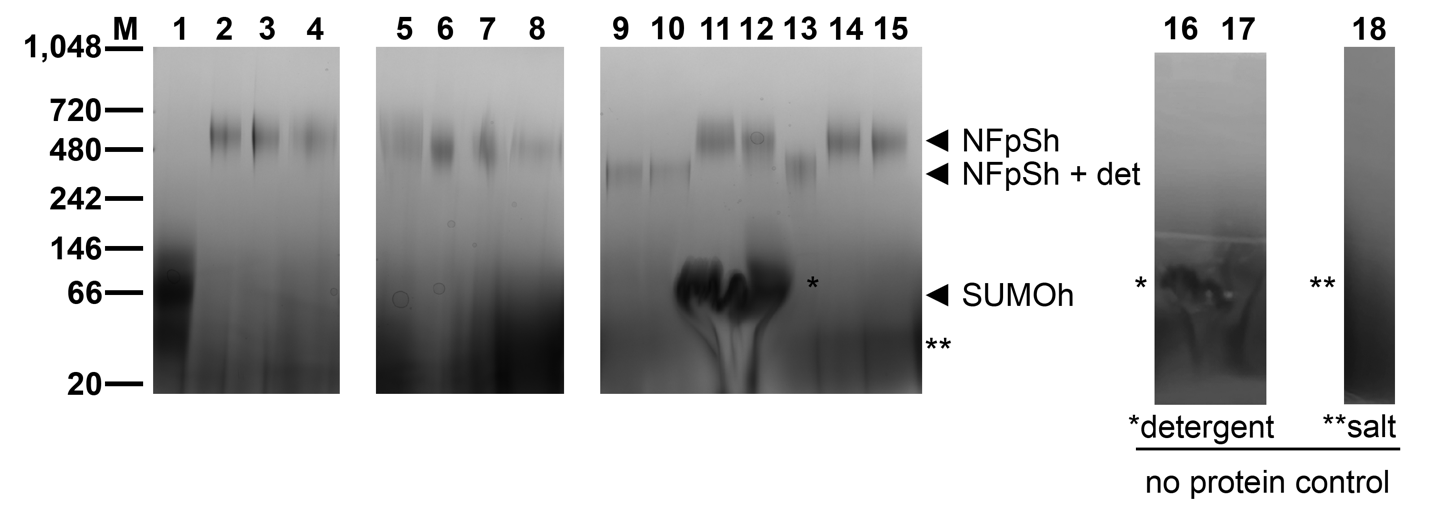

**Supplemental Figure 1: Native-PAGE shows the stability of the N-terminal assembly under a variant of conditions.**

Each lane is as follows:

M – markers (kDa)

1 – SUMO-his

2 – NFPSh (N-terminus of SiRFP with SUMO

3 – NFPSh + 0.5 M urea

4 - NFPSh + 0.5 M guanidine-HCl

5 - NFPSh + 1 M KCl

6 - + NFPSh 1 M urea

7 - NFPSh + 1 M guanidine-HCl

8 - NFPSh + 2 M KCl

9 - NFPSh 1% SDS

10 - NFPSh + 2% SDS

11 - NFPSh + 2% Tween-20

12 - NFPSh + 2% Triton-X

13 - NFPSh + 1% CHAPS

14 - NFPSh + 1 mM FC8

15 - NFPSh + 2 mM FC8

Controls:

16 - 2% Tween-20 only

17 - 2% Triton-X only

18 - 2 M KCl only

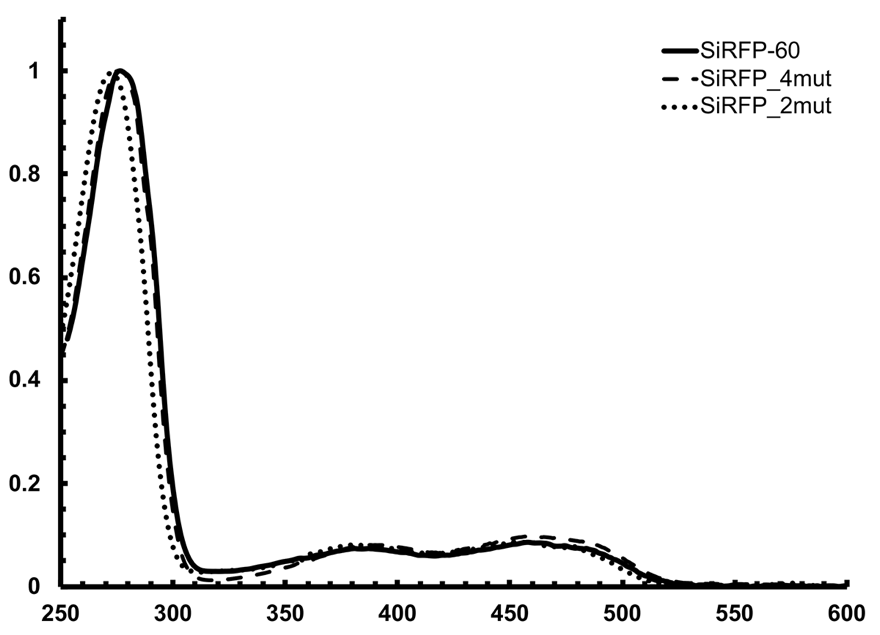

**Supplemental Figure 2: SiRFP_2mut and SiRFP_4mut share spectroscopic signals with SiRFP-60, showing the variants are folded and that flavins bind with full occupancy.**

**Supplemental Figure 3: BN-PAGE analysis of SiRFP variants.** LMW markers (kDa). 1) SiRFP-60 2) SiRHP (64 kDa) 3) SiRFP_2mut 4) SiRFP_4mut 5) SiRFP
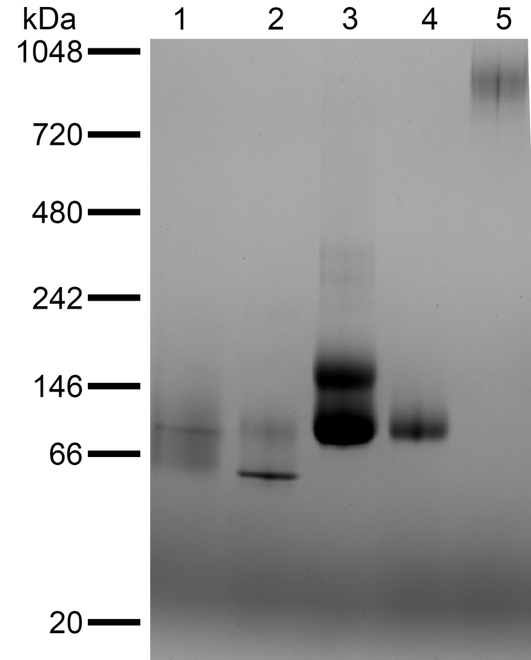
.

**a.**

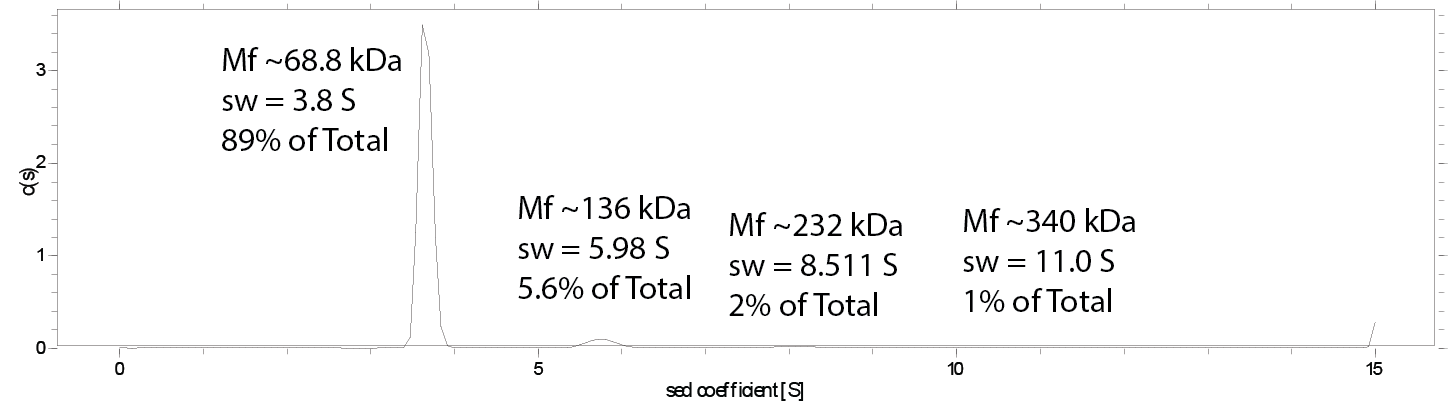
**SiRFP_4mut:**

r.m.s.d. = 0.003808

friction ratio = 1.597

meniscus = 6.023

bottom = 7.1997

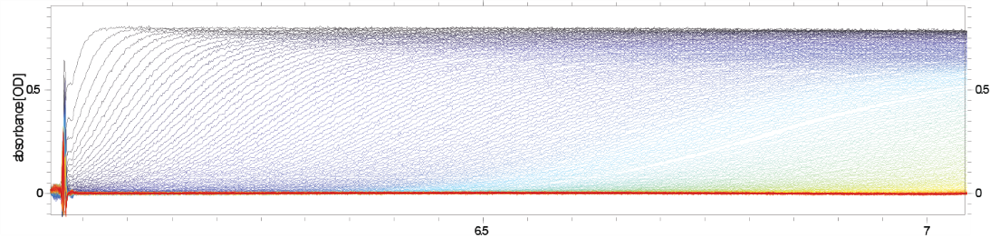

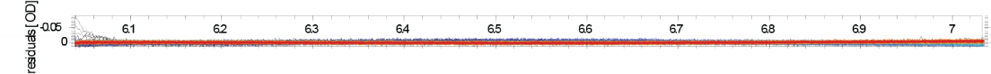

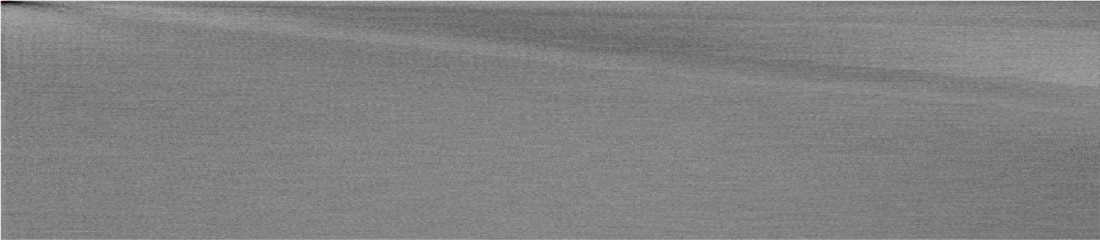

**b.**

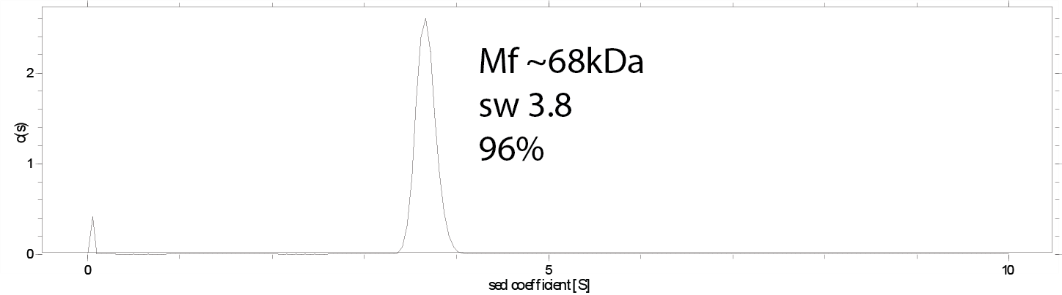
**SiRFP-60:**

r.m.s.d. = 0.003093

friction ratio = 1.576

meniscus = 6.0113

bottom = 7.1997

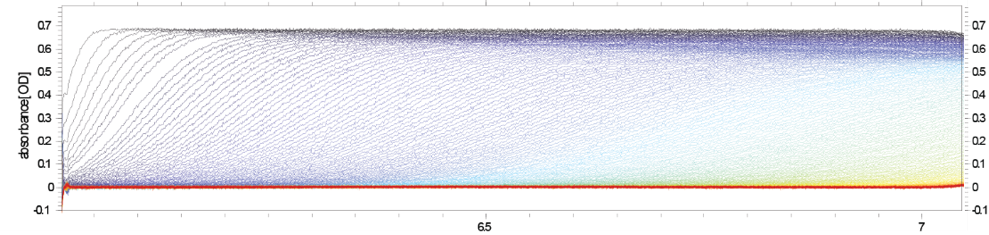

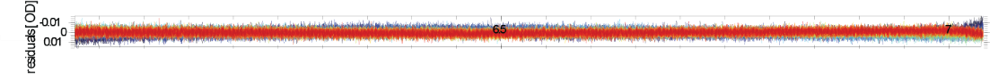

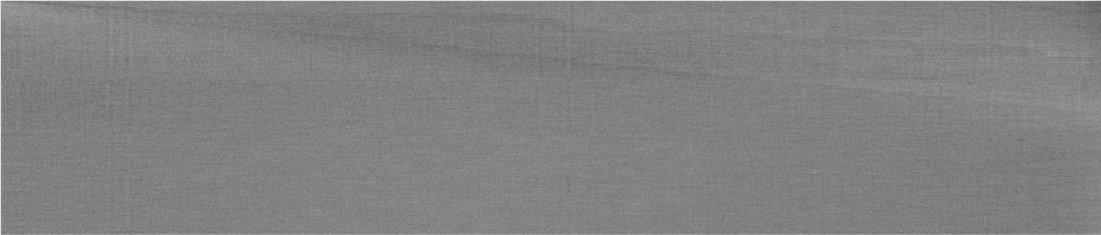

**Supplemental Figure 4: AUC analysis of SiRFP_4mut (a) shows mostly (89%) monomeric SiRFP, with an S value of about 3.8, similar to that of the known monomeric SiRFP-60 (96%) (b).**

**a.**

**SiRFP_4mut:**
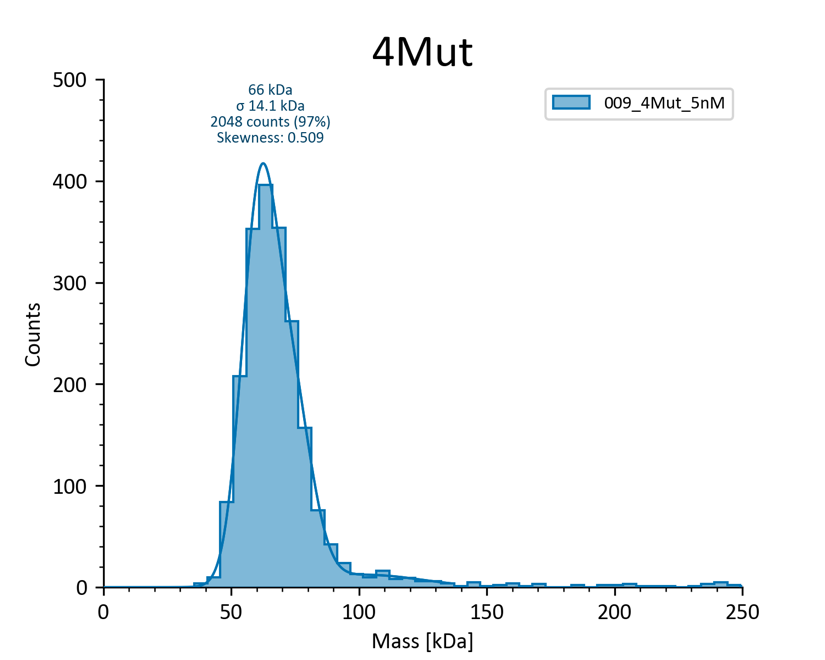

Counts

Mass (kDa)

**b.**

**SiRFP-60:**

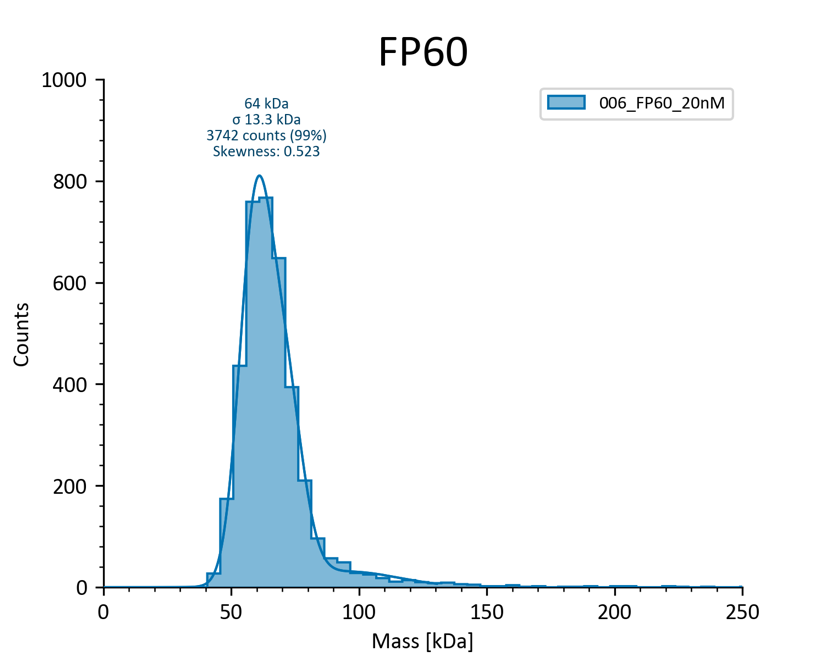

Counts

Mass (kDa)

**Supplemental Figure 5: Mass Photometry. a.** SiRFP_4mut is primarily (97%) a monomer at low (5 nM) concentrations. **b.** SiRFP-60 is also monomeric (99%) even at 4x that concentration (20 nM).

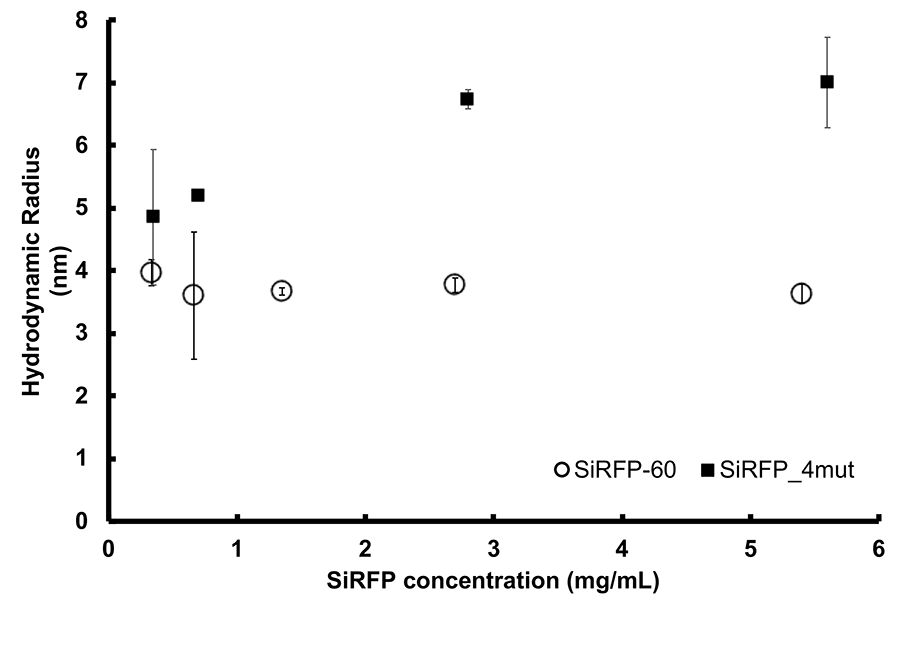

**Supplemental Figure 6:** Dynamic light scattering shows concentration-dependent size of SiRFP_4mut (black squares), in contrast to monomeric SiRFP-60 (circles).

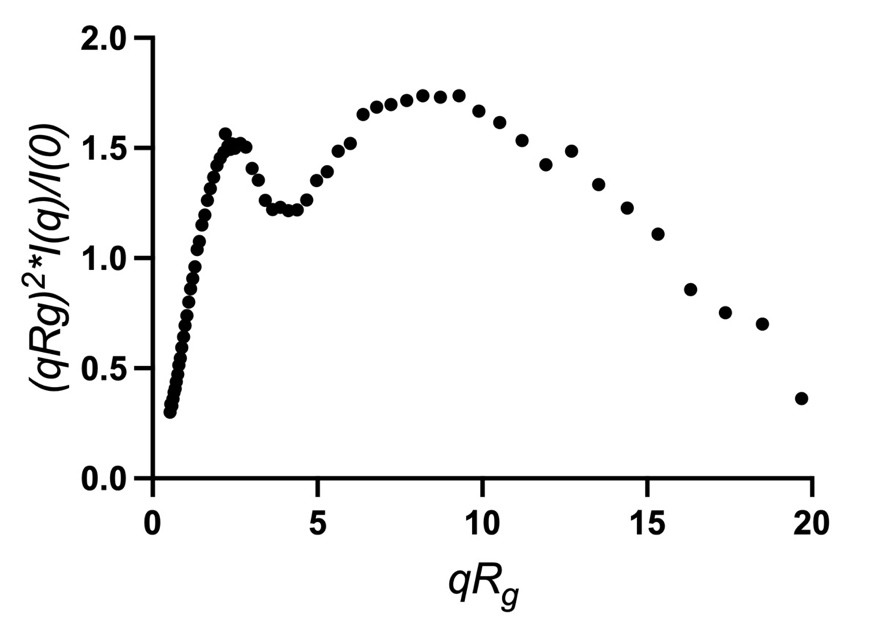

**Supplemental Figure 7:** Dimensionless Kratky analysis shows a flexible tetramer in solution.

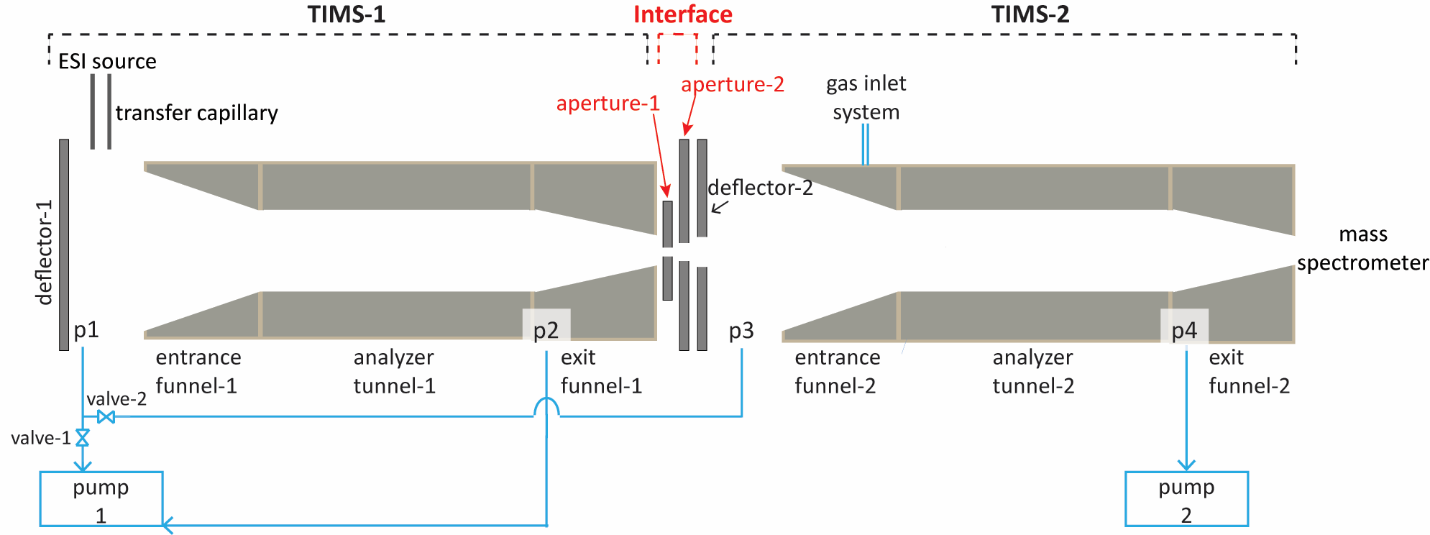

**Supplemental Figure 8:** Detailed schematics of the tandem-trapped ion mobility spectrometer. Reproduced with permission [1].

| **Construct** | **Parent Vector** | **Primers** | **Expression strain** | **Inductant** |
| --- | --- | --- | --- | --- |
| SiRFP_2mut | pBAD | (F) GCCTGGGGCGTACTCAATCAGC | LMG194 | arabinose |
|  |  | (R) AGCGCCAGAAACCCAGGCAAG |  |  |
| SiRFP_4mut | pBAD | synthesized by GenScript | LMG194 | arabinose |
| NFPSh | pET14b | synthesized by GenScript | BL21DE3 | IPTG |

**Table S1: Proteins and expression systems used in this study**

**SUPPLEMENTAL METHODS:**

### **Tandem-TIMS/MS general settings**

The buffer gas was infused into the instrument via the desolvation gas unit of the nano electrospray source with a flow rate of 2.5 L min^-1^. The ion accumulation time for the TIMS1 cartridge was set to 29.83 ms. The desolvation gas temperature was kept at 323 K and the electrospray voltage was set to 875 V. The pressures at p1/p2/p3/p4 (see Figure S2) were kept at 3.1/1.8/0.9/0.4 mbar. RF potentials with a frequency of 566 kHz, 443 kHz, and 3.2 MHz are applied to the TIMS-1 analyzer, TIMS-2 analyzer, and the hexapole, respectively. The RF peak-to-peak amplitude was set to 250 V for both TIMS cartridges. All spectra were acquired in positive ion mode.

### **Tandem-TIMS/MS settings for native measurements**

**TIMS-1:** transmission	**interface:** transmission	**TIMS-2:** analysis

In TIMS-1, a dc bias of 20 V was applied between deflector and entrance funnel, while the dc potential difference across the entrance funnel was kept at 20 V. The dc potentials at analyzer tunnel-1, exit funnel-1, aperture-1, aperture-2, and deflector-2 were set were set to 38/28/28/23/18 V. In TIMS-2, a dc bias of 5V was applied between deflector-2 and entrance funnel-2, and 20 V was kept across the entrance funnel-2. For mobility separation of the WT octamer and the tetramer samples, the potential at the entrance of analyzer tunnel of TIMS-2 was linearly ramped from -7 V to 30 V at a rate of 0.24 V ms^-1^. The dc potential at exit funnel-2 was set to 43 V at all times.

### **Tandem-TIMS/MS settings for activated measurements**

*Mobility selection and collisional activation in the interface of tandem-TIMS*

**TIMS-1**: analysis	**interface**: selection/activation	**TIMS-2**: analysis

In TIMS1, a dc bias of 20 V was applied between deflector and entrance funnel, while the dc potential difference across the entrance funnel was kept at 20 V. For mobility separation, the potential at the entrance of analyzer tunnel of TIMS-1 was linearly ramped from -157 V to -37 V at a rate of 0.76 V ms^-1^. The dc potential at the exit funnel of TIMS-1, aperture-1, aperture-2, and deflector-2 were set to 23/23/18/13 V. In TIMS-2, a dc bias of 5 V was applied between deflector-2 and entrance funnel-2, and 20 V was kept across the entrance funnel-2. For mobility separation in TIMS-2, the potential at the entrance of analyzer tunnel of TIMS-2 was linearly ramped from -12 V to 30 V at a rate of 0.27 V ms^-1^. The dc potential at exit funnel-2 was set to 43 V at all times. To select the WT octamer mobility range, aperture-2 was set to a transmitting voltage (18 V) for a duration of 9.4 ms and at 42.4 ms, and to a blocking potential of 58 V otherwise. For activation experiments, the dc potential at the exit funnel of TIMS-1, aperture-1, aperture-2, and deflector-2 were set to 268/268/263/13 V, with the dc voltage bias set between aperture-2 and deflector-2 of 250 V. All potentials in TIMS-1 analyzer were increased by 245V.
